# Concurrent measures of impulsive action and choice are partially related and differentially modulated by dopamine D_1_- and D_2_-like receptors in a rat model of impulsivity

**DOI:** 10.1101/2022.08.16.504077

**Authors:** Lidia Bellés, Chloé Arrondeau, Ginna Urueña-Méndez, Nathalie Ginovart

**Author notes:** Correspondence: Nathalie Ginovart, Departments of Psychiatry & Basic Neurosciences, Faculty of Medicine, Room E07-2550A, University of Geneva, Rue Michel Servet 1, CH-1211 Geneva, Switzerland. Declarations of interest: none.

## Abstract

Impulsivity is a multidimensional construct, but the relationships between its constructs and their respective underlying dopaminergic underpinnings in the normal population remain unclear. A large cohort of Roman high-(RHA) and low- (RLA) avoidance rats were tested for impulsive action and risky decision-making in the rat gambling task, and then for delay discounting in the delay discounting task to concurrently measure the relationships among the three constructs of impulsivity using a within-subject design. Then, we evaluated the effects of dopaminergic drugs on the three constructs of impulsivity, considering innate differences in impulsive behaviors at baseline. Risky decision-making and delay discounting were positively correlated, indicating that both constructs of impulsive choice are related. Impulsive action positively correlated with risky decision-making but not with delay discounting, suggesting partial overlap between impulsive action and impulsive choice. RHAs showed a more impulsive phenotype in the three constructs of impulsivity compared to RLAs, demonstrating the comorbid nature of impulsivity in a normal population. While amphetamine increased impulsive action and had no effects on risky decision-making regardless of baseline levels of impulsivity, it decreased delay discounting but only in high impulsive RHAs. Conversely, the D_1_R agonist SKF81297, D_3_R agonist PD128907 and D_2/3_R partial agonist aripiprazole decreased impulsive action irrespective of baseline levels of impulsivity, whereas D_2/3_R agonism with quinpirole decreased it exclusively in high impulsive RHAs. Risky decision-making was increased by SKF81297 and quinpirole but not PD128907 and aripiprazole, with quinpirole producing baseline-dependent effects, increasing risky decision-making only in low impulsive RLAs. Finally, while SKF81297, PD128907 and aripiprazole increased delay discounting irrespective of baseline levels of impulsivity, quinpirole decreased it in low impulsive RLAs only. These findings indicate that the acute effects of dopamine drugs were partially overlapping across dimensions of impulsivity, and that only D_2/3_R agonism showed baseline-dependent effects on the three constructs of impulsivity.

## 1. Introduction

Impulsivity is a multidimensional construct that encompasses domains of behavioral inhibition and decision making, and can be broadly categorized into impulsive action (an inability to withhold a motor response) and impulsive choice or decision making (Dalley et al., 2008; Pattij and Vanderschuren, 2008; Winstanley et al., 2006). Impulsive choice involves delay discounting (a preference for small immediate over larger delayed rewards) and risky decision-making (a tendency to prefer disadvantageous or suboptimal options that yield immediate large rewards, despite larger losses in the long-term; Winstanley et al., 2010a). Abnormalities in impulsive action, delay discounting and risky decision-making behaviors have been shown to co-exist in several psychiatric disorders, including substance abuse and addiction (review in Jentsch et al., 2014), attention deficit hyperactivity disorder (Winstanley et al., 2006), and bipolar disorder (Najt et al., 2007) suggesting that the three facets of impulsivity are inter-related and may be mediated by common or overlapping neurobiological mechanisms. To date though, despite a wealth of studies, the relationship between impulsive action, decision-making and risk-taking behaviors in the normal population has produced contrasting evidence as to whether these dimensions are related or not. In humans, some studies show no significant correlation between different components of impulsivity (Broos et al., 2012; McDonald et al., 2003; Reynolds et al., 2008), whereas others show correlations between action and choice dimensions of impulsivity (Dougherty et al., 2009). Studies in animals have also produced mixed results. A number of studies showed that impulsive action is unrelated to delay discounting (Broos et al., 2012; Winstanley et al., 2004) and to risky decision-making (El Massioui et al., 2016; Rivalan et al., 2013), and that delay discounting and risky decision-making are also unrelated (Freels et al., 2020; Mitchell et al., 2012), suggesting not only that impulsive choice and action are independent behaviors but also that even sub-dimensions of impulsive choice are dissociable. On the other hand, there is also evidence for positive associations between impulsive action and delay discounting (Moreno et al., 2010; Robinson et al., 2009), or risky decision-making (Barrus et al., 2015; Higgins et al., 2018; Tremblay et al., 2021), and between delay discounting and risky decision-making (Kirkpatrick et al., 2014). Inconsistencies may stem from a number of factors, including methodological differences between studies such as the use of different behavioral tasks to measure the same construct of impulsivity. For instance, in rodents, the risky decision-making task (RDT), the probability discounting task (PDT), and the rat gambling task (rGT) are used interchangeably to measure risky decision-making, although those tasks are likely tapping slightly different processes (review in Yates, 2019). Similarly, the Go/NoGo or the Five-choice serial-reaction time task (5-CSRTT) are used as measures of impulsive action although evidence indicate that the two tasks may actually be measuring different aspects of impulsive action, namely “action cancelation” on the Go/NoGo and “action restraint” on the 5-CSRTT (review in Dalley et al., 2011). Alternatively, inconsistencies may also be due to variations in environmental conditions among studies, which are known to influence impulsive behaviors (Belles et al., 2021; Zeeb et al., 2013). It is also worth noting that the large majority of work in this area did not perform within-subject correlations but rather compared the associations between indices of impulsivity at the population level and no studies in rodents have examined more than two behavioral measures of impulsivity in a same cohort of animals. Moreover, it can reasonably be expected that the narrow range and small inter-individual variations in impulsivity found in inbred rat strains may hamper the detection of possible relationships between behavioral indices of impulsivity. In that respect, expanding the range of impulsivity using subjects with varying degrees of innate impulsivity might help to improve our understanding of their precise relationships to one another. Altogether, the existence of conflicting results in the field, the limited number of studies using within-subject approaches and the lack of studies investigating concurrently the three facets of impulsivity in a same cohort of animals, emphasize the need for more systematic investigations to understand more comprehensively the relationships between behavioral indices of impulsivity.

A wealth of studies supports a central role of dopamine (DA) in all facets of impulsivity (review in Dalley and Roiser, 2012; Yates, 2019), potentially through abnormal modulation of DA receptor function in striatum. Indeed, rats showing high impulsive action (Belles et al., 2020; Dalley et al., 2007) or high delay discounting (Barlow et al., 2018b) exhibit reduced densities of striatal DA D_2/3_ receptors (D_2/3_R). However, pharmacological manipulations of DA receptor subtypes on the different facets of impulsivity have produced mixed results. Specifically, systemic administration of D_2/3_R agonists have been reported to either decrease (Fernando et al., 2012; Winstanley et al., 2010b) or have no effect (Zeeb et al., 2009) on impulsive action, and to either increase (Georgiou et al., 2018; Wallin-Miller et al., 2018), decrease (Simon et al., 2011; Smith et al., 2018) or have no effect (Swintosky et al., 2021; Zeeb et al., 2009) on risky decision-making. Similarly, selective D_3_R agonists produced either an increase (Barrus and Winstanley, 2016), a decrease (Barrus and Winstanley, 2016; St Onge and Floresco, 2009) or have no effect (Di Ciano et al., 2015; Swintosky et al., 2021) on impulsive action and risky decision-making, whereas selective D_1_R agonists increase (Zhu et al., 2017), decrease (Pekcec et al., 2018; Winstanley et al., 2010b), or have no effect (Zeeb et al., 2009) on impulsive action, and increase (St Onge and Floresco, 2009; Zeeb et al., 2009) or have no effect on risky decision-making (Oinio et al., 2017; Wallin-Miller et al., 2018). There are multiple sources from which these inconsistencies could arise, including, here again, methodological differences between studies. However, the majority of studies investigating the effects of DA agents on impulsivity did not consider baseline differences in impulsive behavior, which may significantly affect the response to drug challenges and could thus also be a source of inconsistencies. For instance, although relatively scare, previous findings indicate that the effect of psychostimulants on behavioral inhibition is baseline-dependent, reducing impulsivity in animals that exhibit high impulsive behavior at baseline, and increasing it or having no effect in animals displaying low levels of baseline impulsivity (Caprioli et al., 2015; Feola et al., 2000; Tomlinson et al., 2014), although contrasting results have also been reported (Barbelivien et al., 2008; Higgins et al., 2021). This raises the important point that variations in baseline levels of impulsive behavior can produce distinct outcomes and even prevent the detection of effects and emphasizes the need to consider basal levels of impulsivity when investigating drug effects.

In this context, an interesting preclinical model of impulsivity is the Roman rat lines, which display a wide range of impulsivity levels, with Roman high-avoidance (RHA) rats exhibiting higher levels of impulsive action and delay discounting compared to Roman low-avoidance (RLA) rats (Belles et al., 2020; Moreno et al., 2010). Although no data are yet available regarding the risky-taking profile of these two rat lines, the impulsive phenotype in RHAs is related to lower levels of striatal D_2/3_R availabilities, to increased levels of amphetamine (AMPH)-induced DA release and to a greater propensity to cocaine abuse (Belles et al., 2020; Dimiziani et al., 2019). As current evidence strongly implicates abnormalities in DA neurotransmission in both impulsive action and impulsive choice (review in Dalley and Ersche, 2019; Dalley and Roiser, 2012), the divergent impulsive and neurochemical profiles expressed by the Roman lines make them an interesting model to investigate the interaction between different facets of impulsivity and examine potential differential effects of DAergic drugs on impulsive behaviors. The aim of the present study was two-fold: 1) to concurrently measure, using a within-subject repeated measure design, the relationships among impulsive action, risky decision-making and delay discounting, and 2) to further examine the effects of dopaminergic drugs on the three facets of impulsivity, taking into consideration innate differences in impulsive behaviors at baseline. To these aims, a large cohort of RHA and RLA rats were first tested for motor impulsivity and risky decision-making in the rGT, and then for delay discounting in the delay discounting task (DDT). After each task, innate baseline impulsive behavior was established and the effects of DA agonist and antagonists on the animals’ performance in both tasks were evaluated.

## 2. Material and methods

### 2.1. Animals

Three-month-old male RHA (n=24) and RLA (n=24) rats from our permanent colony of outbred Roman rats at the University of Geneva were used. Animals were housed three per cage and maintained under a 12-hour-light-dark cycle (lights on at 7:00h), with controlled temperature (22 ± 2◦C) and humidity (50–70%). Rats were food deprived and maintained at 85% to 90% of their free-feeding weight, whereas water was provided ad libitum. All experimental procedures were approved by the Animal Ethics Committee of the canton of Geneva, and husbandry was performed in accordance with the Swiss Federal Law on animal care.

### 2.2. Experimental timeline

Rats were first trained and tested for motor impulsivity and risky decision-making in the rat Gambling Task (rGT). Once stable baseline behavior measures were established, the effects of the DA releaser D-Amphetamine (AMPH), the D_3_R agonist PD128907, the D_2/3_R partial agonist aripiprazole and the D_2_R antagonist L-741,626 was measured in the rGT. In each rat line, each rat was pseudo-randomly assigned for testing with three drugs only in a counterbalanced order (n=12 rats per drug in each line). Subsequently, the same cohort of animals were trained and tested for delay discounting in the delay discounting task (DDT). Once stable baseline behavioral measures were established, animals were tested in the DDT using the same pharmacological challenges as those used in the rGT.

### 2.3. Drugs and doses

Drug doses and pretreatment times are provided in Table S1. All doses were calculated as the salt. Aripiprazole (1.0 mg/kg), L-741,626 (2.0 mg/kg), PD128907 (0.3 mg/kg), quinpirole (0.5 mg/kg) and SKF81297 (0.5 mg/kg) were purchased from Tocris Bioscience (Bristol, UK). Amphetamine (AMPH; 1.0 mg/kg) was purchased from Sigma-Aldrich (Buchs, Switzerland). AMPH, SKF81297, quinpirole and PD128907 were dissolved in 0.9% sterile saline. Aripiprazole and L741,626 were dissolved in 5% tween 80 and diluted in distilled water. All drugs were freshly prepared and injected intraperitonially (i.p.) in a volume of 1 ml/kg either 20 min (aripiprazole, PD128907), 30 min (SKF81297), 60 min (L741,626), or 90 min (AMPH, quinpirole) before testing. Doses and routes of administration were based on previous reports (Besson et al., 2010; Blaes et al., 2018; Di Ciano et al., 2015; St Onge and Floresco, 2009; Tournier et al., 2013; Zeeb et al., 2009).

### 2.4. Rat Gambling Task (rGT)

Rats were trained on the rGT using eleven modular operant chambers each encased within a ventilated sound-attenuating cubicle (Med Associates Inc., St. Albans, VT). Each chamber had a houselight and was equipped with a curve wall including 5 equally spaced nose-poke apertures, each provided with a light inside. The opposite wall was equipped with a pellet receptacle with a light and connected to an external dispenser. Only the outer four of the five holes were used (i.e., the central hole was inactivated).

Details on the rGT procedure are provided in the SOM. All rats were trained on the rGT, as described previously (Zeeb et al., 2009). Briefly, rats were trained to nose poke in one of the four holes that was illuminated to receive a food pellet reward (Carli et al., 1983). Animals were then trained on a forced-choice version of the rGT, where only one of the four possible options was presented on each trial, to ensure equal experience with all the four reinforcement contingencies and to avoid preferences toward a particular hole. Rats were then moved to the rGT in which all options were simultaneously available during each trial. A trial started by a nose poke into the illuminated food tray. After an inter-trial interval (ITI) of 5 sec, four stimulus lights were turned on and the animal required to nose poke into one of these holes within 10 sec. This response was rewarded or punished depending on the reinforcement contingency for that hole option (Table S2). The optimal choice in rGT was P2, as this option is the most rewarded per unit time. The next best option was P1, and the two disadvantageous options were P3 and P4, due to the lower probability of receiving reward and the longer punishing TO periods incurred (Table S2). Missed trials (i.e., omissions) were also recorded and all stimulus lights were extinguished, and the food tray illuminated, allowing the animal to start another trial. Responses made before the onset of the visual stimulus (i.e., during the ITI) were considered as premature or impulsive and resulted in a 5-sec TO period, after which the food tray light was turned on and animals could initiate the next trial. Two versions of the rGT were used which differed only in the spatial location of the options and were counterbalanced across all animals (Table S2). Animals were tested until they acquired stable response, then baseline measures were established across three consecutive sessions (<10% variation in each response hole). The outcome measures were the percentage of premature responses [(#premature responses/#total number of trials initiated)*100] as an index of motor impulsivity; choice score [(P1+P2)–(P3+P4)*100] as a measure of risky decision-making in rats (Adams et al., 2017; Ferland and Winstanley, 2017); the percentage of omissions [(#omission responses/#total number of trials initiated)*100]; the total number of trials completed; the choice latency (in seconds) which was the time from the end of the ITI to a nose-poke response into a hole and the collect latency (in seconds) which was the time from reward delivery to reward collection into the food tray. The effect of pharmacological manipulations was then tested. All drugs were prepared fresh daily, and the order of administration was counterbalanced. Each rat was tested with only three drugs (12 RHA and 12 RLA rats per drug) in a 2-day cycle, in which animal received i.p. vehicle (day 1) and then drug (day 2) injections. To prevent any potential carryover effects, animals were given a washout period between drug of at least 3 days.

### 2.5. Delay Discounting Task (DDT)

Rats were trained in the DDT using eleven modular operant chambers each encased within a ventilated sound-attenuating cubicle (Med Associates Inc., St. Albans, VT). Each chamber was equipped with a houselight, two nose-poke holes and a food tray with a light. The two nose-poke holes were positioned on the left and on the right side of a lateral wall with the food tray in-between and were each equipped with a cue-light.

Details on the DDT procedure are provided in the SOM. The same cohort of rats were first trained for magnitude discrimination using a modified version of the amount-discrimination training (Renee Renda et al., 2018). The session began in darkness and trials started at 72-sec inter-trial interval. Each trial began with the illumination of the food tray. Animals were trained to make a nose poke response in the food tray, ensuring that they were centrally located at the start of the trial. Upon a successful nose poke, the food tray light was extinguished and one (forced-choice trials) or the two (free-choice trials) holes were illuminated. A response in the small reinforcer hole resulted in the delivery of 1 pellet, whereas a response on the larger reinforcer hole resulted in the delivery of 3 pellets. There was no delay in the delivery of either the small or large reinforcers in this phase. Rats were required to reach a criterion of 60 completed trials with ≥80% preference for the larger reinforcer across three consecutive sessions. Animals were then moved to the DDT (Barlow et al., 2018b; Isherwood et al., 2017). Each session was divided into 6 blocks of 12 trials, constituted of 4 forced-choice trials followed by 6 free-choice trials. Similar to the training sessions, the task began in darkness and trials started at 72-sec inter-trial interval. Each trial began with the illumination of the food tray. If rats nose poked, the food tray light extinguished following the illumination of one (forced-choice trials) or the two (free-choice trials) holes. One hole was designated as the delay hole and the other as the immediate hole. A nose-poke into the immediate hole turned on the food tray light and one pellet was delivered with no time delay (i.e., 0 sec). If the choice was the delay hole, the stimulus light within the hole remained turned on for the duration of the delay. Following the delay, the stimulus light was switched off, the food tray light turned on and 3 pellets were delivered. Delays were increased across blocks from 0 to 2, 4, 8, 16 and 32 sec. An adjusting inter-trial interval ensured that a new trial started every 72 sec. A failure to respond either on the food tray or on one of the holes within 10 sec (i.e., omissions) resulted in the extinction of the light and the inter-trial interval initiated before the next trial. Animals were tested until they acquired stable response, then baseline measures were established across three consecutive sessions (<10% variation in the delay hole responses). Delay-discounting curves were obtained by plotting the percentage choice for the large reward as a function of delay (sec). The area under the curve (AUC) was used as a measure of delay discounting (Magnard et al., 2018; Myerson et al., 2001). The AUC values ranged from 0 to 1, with lower AUC values indicating greater delay discounting, and thus higher levels of delay-related impulsivity. Choice latency (in seconds) was calculated measured the time from the end of the ITI to a nose-poke response into a hole. Collect latency (in seconds) measured the time from reward delivery to reward collection into the food tray. Pharmacological manipulations were then begun. Rats were tested with the same drugs, doses, and pretreatment times as those used for the rGT.

### 2.6. Data analyses

Choice score data were arcsine transformed to minimize the effect of an artificially imposed ceiling. Distribution of the data was tested for normality with the Shapiro-Wilk test. Non-normal data were normalized using log-10 transformation. Between-line differences in percentage of premature responses, percentage of omissions, choice score, AUC, total trials completed, choice and collect latency at baseline were analyzed with a 2-tailed independent Student’s *t* test. Between-line differences in delay-discounting curves were analyzed with a two-way repeated measures ANOVA with delay as within-subject factor and line as between-subject factor.

The effects of pharmacological challenges on premature responding, choice score, AUC, omissions, latencies and total trials completed were analyzed using a two-way mixed factorial ANOVA with line (i.e., RHA and RLA) and treatment as within-subject factors. Post-hoc comparisons were performed using paired *t* test when comparing data within-line, and independent *t* test when comparing data between-line. Correlations between motor impulsivity, risky decision-making, delay discounting, choice and collect latency were performed using Pearson’s correlation coefficient. All statistical analyses were carried out using SPSS Statistics 27.0 software (IBM Corp., Armonk, NY, USA). The significance level was set at *P*<0.05.

## 3. Results

### 3.1. Baseline performance in the rGT and DDT

Consistent with previous reports using the 5-CSRTT (Belles et al., 2020; Moreno et al., 2010), when assessed using the rGT, RHA rats made significantly more premature responses than RLAs (Fig.1A; *t*=7.53, *P*<0.001), indicating greater motor impulsivity in the former line.

**Fig. 1:**
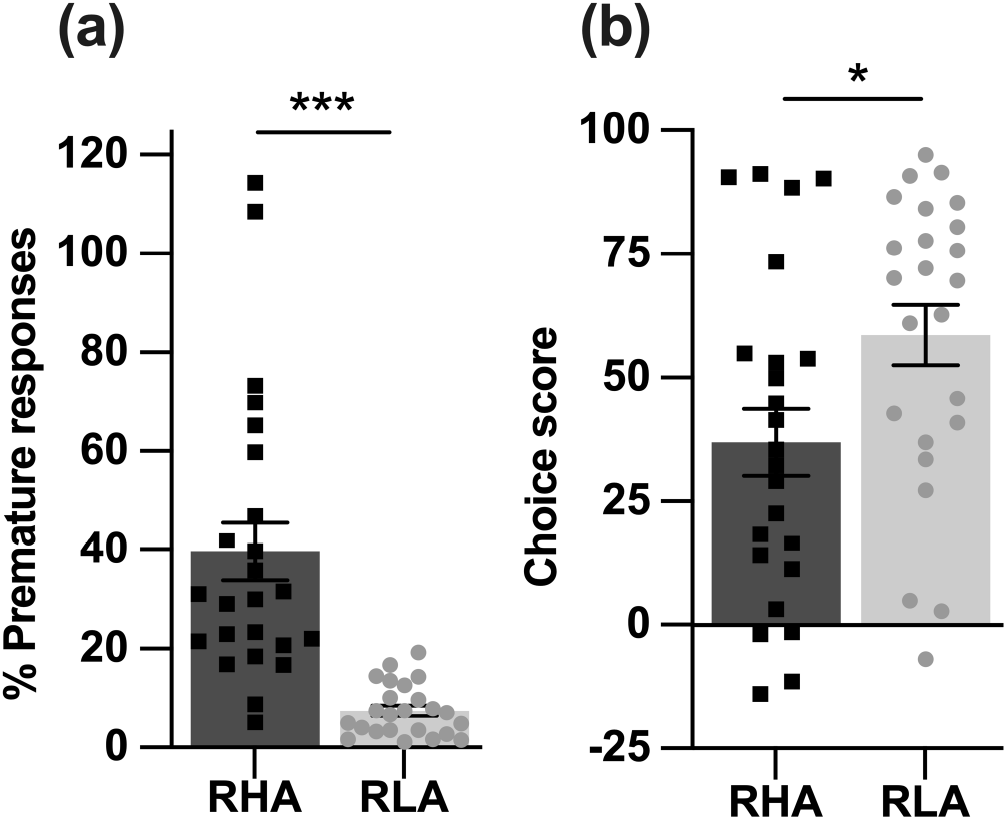
Performance of RHA (n= 24) and RLA (n= 24) rats on the rGT. (a) Percentage of premature responses. (b) Choice score. Data are expressed as mean±SEM. *P<0.05 and ***P<0.001 using a 2-tailed independent Student’s t test.

RHA rats also exhibited lower choice score than RLAs in the rGT (Fig. 1B; *t*=2.30, *P*=0.03), indicating that high impulsive RHA rats also displayed greater risky decision-making. RHA rats were significantly faster than RLA rats to select a response in the rGT, as indicated by a shorter response latency (Table 1; *t*=-4.90, *P*<0.001), but both lines did not differ in the latency to collect the reward (Table 1; *t*=1.10, *P*=0.29).

**Table 1.** Choice and collect latencies of RHA and RLA rats on the rGT and DDT.

| Group | rGT |  | DDT |  |
| --- | --- | --- | --- | --- |
|  | Choice latency (s) | Collect latency (s) | Choice latency (s) | Collect latency (s) |
| RHA | **1.70±0.16\*\*\*** | 1.02±0.03 | **1.48±0.06\*\*\*** | **0.48±0.02\*** |
| RLA | 2.61±0.09 | 0.97±0.04 | 1.87±0.07 | 0.43±0.03 |

Data are mean ± SEM. \* $P<0.05$ and \*\*\* $P<0.001$ using 2-tailed independent Student's $t$ test.

**Table 2.** Relationships between choice and collect latencies and premature responses, choice score and AUC on the rGT and DDT.

|  |  |  | Choice latency | Collect latency |
| --- | --- | --- | --- | --- |
| rGT | %Premature responses | Pearson's $r$ | **-0.78\*\*\*** | 0.12 |
| | | $P$-value | 0.001 | 0.42 |
| | Choice score | Pearson's $r$ | **0.39\*\*** | 0.16 |
| | | $P$-value | 0.006 | 0.27 |
| DDT | AUC | Pearson's $r$ | **0.37\*\*** | **-0.53\*\*\*** |
| | | $P$-value | 0.01 | 0.001 |

\*\* . Correlation is significant at the 0.01 level.
\*\*\* . Correlation is significant at the 0.001 level.

A comparison of the mean delay discounting curves obtained at baseline in RHA and RLA rats is shown in Fig. 2A.

**Fig. 2:**
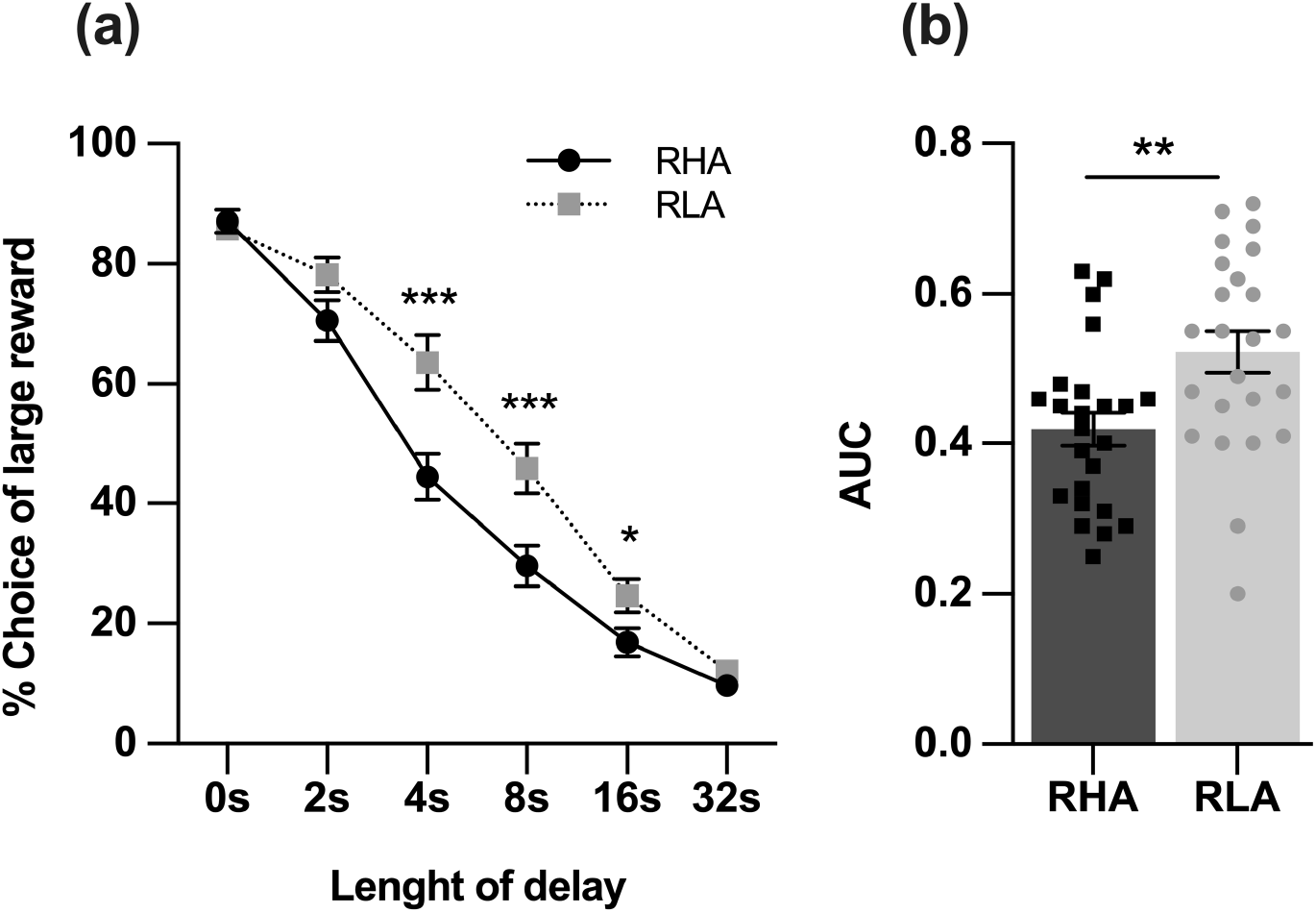
Performance of RHA (n= 24) and RLA (n= 24) rats on the DDT. (a) Delay discounting curves obtained in both lines by plotting the percentage choice of large reward as a function of length of delay (sec). (b) AUC. Data are expressed as mean±SEM. **P<0.01 and ***P<0.001 using (a) two-way repeated measures ANOVA and (b) 2-tailed independent Student’s t test.

At the 4-, 8- and 16-sec delays, RHAs showed more preference for the small, immediate reward vs. large, delayed rewards when compared to RLA rats (Fig. 2A; line: *F*_1,46_=7.57, *P*<0.01; delay: *F*_5,230_=354.20, *P*<0.001; line x delay: *F*_5,230_=6.15, *P*<0.001). Consistent with this, RHAs showed lower AUC than RLAs (Fig. 2B; *t*=2.92, *P*<0.01), indicating greater delay discounting in the former line. Similar to the rGT, RHA rats displayed shorter choice latency in the DDT (Table 1; *t*=-4.17, *P*<0.001), but showed a longer collect latency in comparison to RLA rats (Table 1; *t*=1.98, *P*=0.05).

### 3.2. Relationships between premature responding, choice score and AUC at baseline

When RHAs and RLAs data were combined for analyses, premature responding was negatively correlated with choice score in the rGT (Fig. 3A; *r*=-0.42, *P*=0.003), but did not correlate with AUC in the DDT (Fig. 3B; *r* = −0.21, *P*=0.16). Additionally, choice score was positively correlated with AUC (Fig. 3C; *r*=0.39, *P*=0.006).

**Fig. 3:**
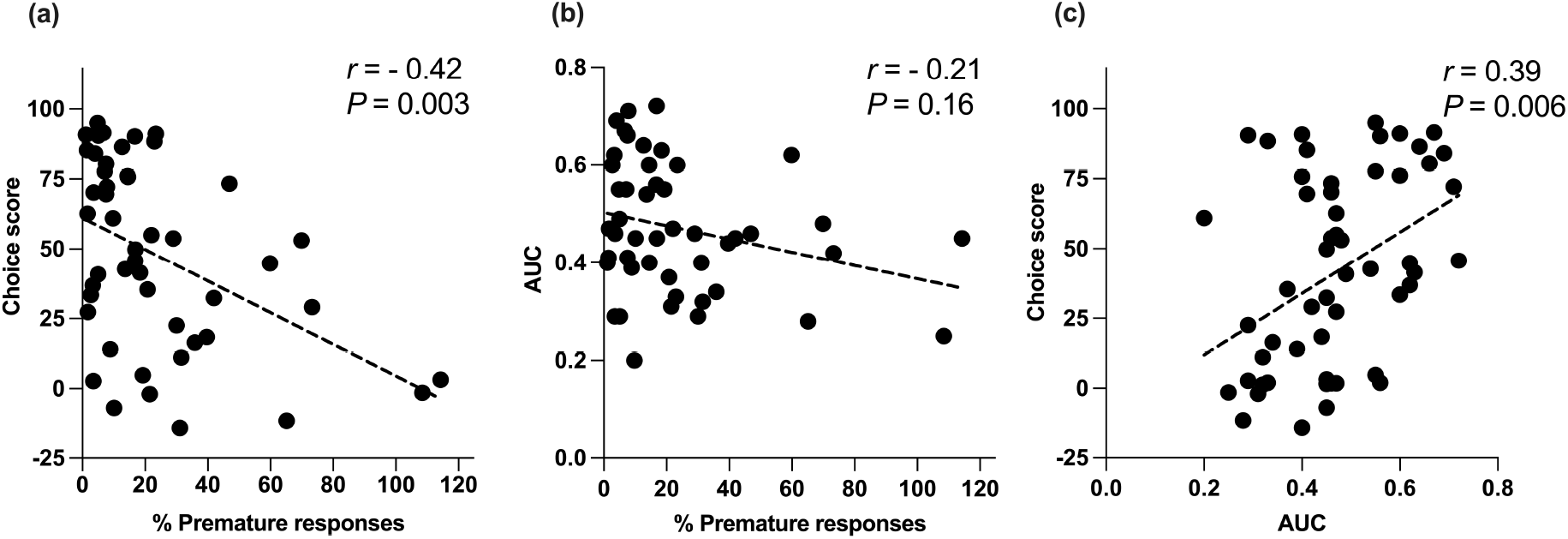
Relationships between impulsive action, risky decision-making, and delay discounting. While (a) premature responses were negatively correlated with choice score, (b) no correlation was found between premature responses and AUC. (c) Additionally, choice score was positively correlated with AUC. Correlations were tested using Pearson’s correlation coefficient (n= 48).

Interestingly, choice latency in the rGT negatively correlated with premature responding (*r*=-0.78, *P*<0.001), whereas it positively correlated with choice score (*r*=0.39, *P*=0.006). In the DDT, choice latency positively correlated with AUC (*r*=0.37, *P*=0.01). These results suggest that in both tasks, speed of responding was predictive of high impulsive behavior. Unlike choice latency, there was no significant correlation between collect latency and premature responses (*r*=0.12, *P*=0.42) or choice score in the rGT (*r*=0.16, *P*=0.27). However, collect latency was negatively correlated with AUC in the DDT (*r*=-0.53, *P*<0.001), indicating that motivation for food reward was predictive of low delay-related impulsive choice.

### 3.3. Dopaminergic modulation of premature responding

The effects of drugs acting on DA transmission on premature responding in high impulsive RHA and low impulsive RLA rats are shown in Fig. 4. Two-way mixed factorial ANOVA showed an effect of treatment (*F*_11,209_=23.66, *P*<0.001), line (*F*_1,19_=16.14, *P*<0.001) but no treatment x line interaction (*F*_11,209_=1.70, *P*=0.07) on premature responding. Post hoc contrasts indicated that AMPH significantly increased premature responding in both RHA (*P*<0.001) and RLA (*P*<0.001) rats (Fig. 4), with no significant changes in omitted trials, in choice latency, or in collect latency (all *P*>0.05; Table S3). In contrast, premature responding was significantly decreased in both lines following treatments with the direct acting D_1_R agonist SKF81297 (RHA: *P*<0.001; RLA: *P*=0.05), and D_3_R agonist PD128907 (RHA: *P*=0.005; RLA: *P*=0.01; Fig. 4). PD128907, but not SKF81297, had significant effects on omitted trials (*P*<0.01) and collect latency (*P*<0.05) in RHA but not in RLA rats (Table S3), with no effect on the choice latency, which may reflect a decreased motivation rather than a general decrease in motor output in RHA rats. Interestingly, a postsynaptic dose of the D_2/3_R agonist quinpirole (i.e., 0.5 mg/kg; Eilam and Szechtman, 1989; Horvitz et al., 2001) decreased premature responding in RHAs (*P*=0.004) but not in RLAs (*P*=0.99). A notable effect of quinpirole was an increase in omitted trials (*F*_11,209_=14.23, *P*<0.001; Table S3), which was significant for both RHA and RLA rats (RHA: saline vs quinpirole: *P*<0.001; RLA: saline vs quinpirole: *P*<0.05). Similarly, there were significant effects of quinpirole on choice and collect latencies (*F*_7,154_=27.38, *P*<0.001; Table S3), which were both significantly slowed in RHAs (choice latency: *P*<0.01; collect latency: *P*<0.05) but not in RLAs (choice latency: *P* >0.05; collect latency: *P* <0.01). The partial D_2/3_R agonist aripiprazole decreased premature responding in both lines (RHA: *P*=0.01; RLA: *P*=0.03), with no significant changes in omitted trials, in choice or collect latencies (all *P*>0.05; Table S3). Blocking D_2_R with L-741,626 had no effect on premature responding (all *P*>0.05; Fig. 4),, omitted trials, or in choice and collect latencies (all *P*>0.05; Table S3) in either RHAs or RLAs.

**Fig. 4:**
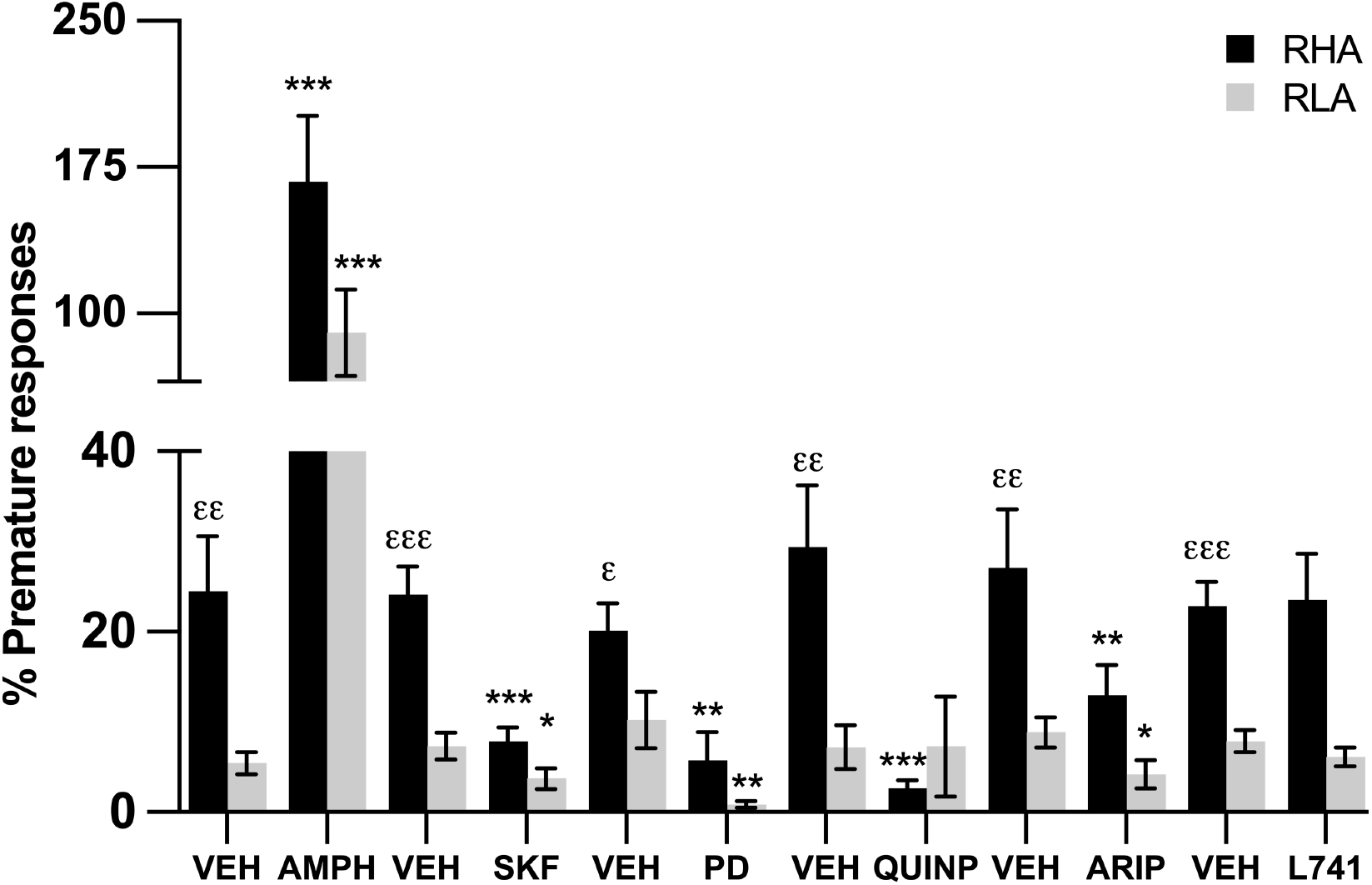
Dopaminergic modulation of premature responding in RHA (n= 11-12) and RLA (n= 10-12) rats. Pharmacological manipulations were performed with i.p. injections of: vehicle (VEH), amphetamine (AMPH; 1.0 mg/kg; i.p.), SKF81297 (SKF; 0.5 mg/kg; i.p.), PD128907 (0.3 mg/kg; i.p.), quinpirole (QUINP; 0.5 mg/kg; i.p.), aripiprazole (ARIP; 1.0 mg/kg; i.p.) and L741,626 (L741; 2.0 mg/kg; i.p.). Drug presentation was counterbalanced across animals. Data are expressed as mean±SEM. *P<0.05, **P<0.01 and ***P<0.001 significantly different from the respective vehicle; εεP<0.01 and εεεP<0.001 significantly different from RLA vehicle using a two-way mixed factorial ANOVA.

### 3.4. Dopaminergic modulation of risky decision-making

The effects of drugs acting on DA transmission on choice score depending on the rat line are shown in Fig. 5. When comparing the RHA and RLA lines, a main effect of treatment (*F*_11,209_=1.91, *P*=0.04) but no main effect of line (*F*_1,19_=0.27, *P*=0.61), and no treatment x line interaction (*F*_11,209_=0.63, *P*=0.80) were found. Post-hoc contrasts showed that D_1_R stimulation with SKF81297 decreased choice score in both RHA (*P*=0.04) and RLA rats (*P*<0.001). Interestingly, the D_2/3_R agonist quinpirole decreased choice score in RLA rats (*P*=0.007) but not in RHA rats (*P*=0.54). None of the other DA acting drugs used produced significant changes in risky decision-making in either RHAs or RLAs (Fig. 5).

**Fig. 5:**
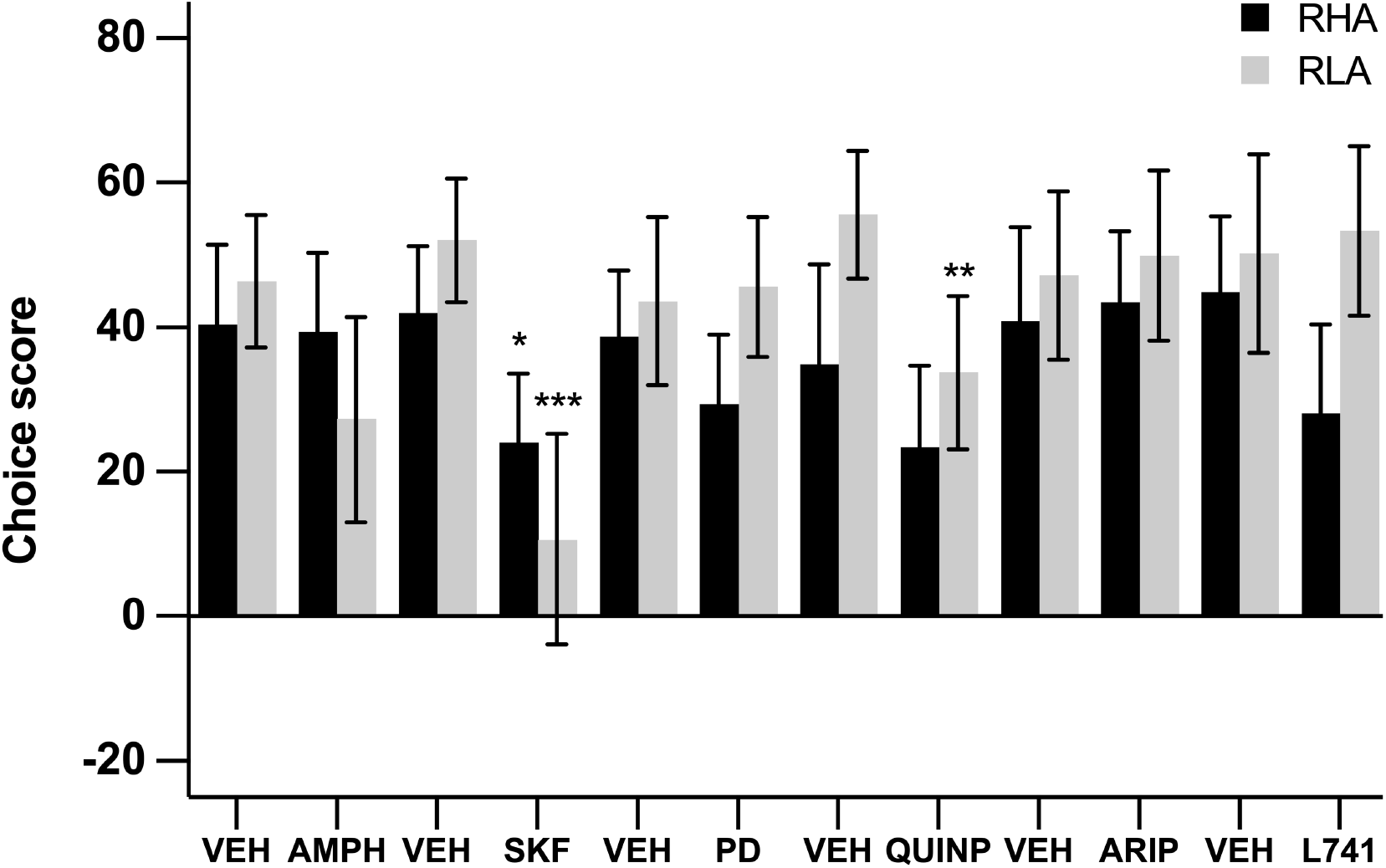
Dopaminergic modulation of choice score in RHA (n= 11-12) and RLA (n= 10-12) rats. Pharmacological manipulations were performed with i.p. injections of: vehicle (VEH), amphetamine (AMPH; 1.0 mg/kg; i.p.), SKF81297 (SKF; 0.5 mg/kg; i.p.), PD128907 (0.3 mg/kg; i.p.), quinpirole (QUINP; 0.5 mg/kg; i.p.), aripiprazole (ARIP; 1.0 mg/kg; i.p.) and L741,626 (L741; 2.0 mg/kg; i.p.). Drug presentation was counterbalanced across animals. Data are expressed as mean±SEM. *P<0.05, **P<0.01 and ***P<0.001 significantly different from the respective vehicle using a two-way mixed factorial ANOVA.

### 3.5. Dopaminergic modulation of delay discounting

Two-way mixed factorial ANOVA showed an effect of treatment (*F*_11,209_=14.56, *P*<0.001), line (*F*_1,19_=5.63, *P*=0.03) and an interaction of treatment x line (*F*_11,209_=1.85, *P*=0.05) on AUC as an index of delay discounting (Fig. 6). Post hoc contrasts indicated that AMPH increased AUC in RHA (*P*=0.01) but not in RLA (*P*=0.90) rats. In contrast, D_1_R agonist stimulation with SKF81297 decreased AUC in RHA (*P*=0.008) and tended to decrease it in RLA (*P*=0.06) rats, whereas D_3_R stimulation with PD128907 significantly decreased AUC in both rat lines (RHA: *P*=0.03; RLA: *P*=0.001). Interestingly, D_2/3_R stimulation with quinpirole increased AUC in RLA (*P*=0.05) but not in RHA (*P*=0.87) rats. Finally, in both RHA (*P*<0.001) and RLA (*P*=0.009) rats, AUC was decreased by aripiprazole. Blockade of D_2_R with L-741 had no effect on AUC in either RHAs (*P*=0.09) or RLAs (*P*=0.06).

**Fig. 6:**
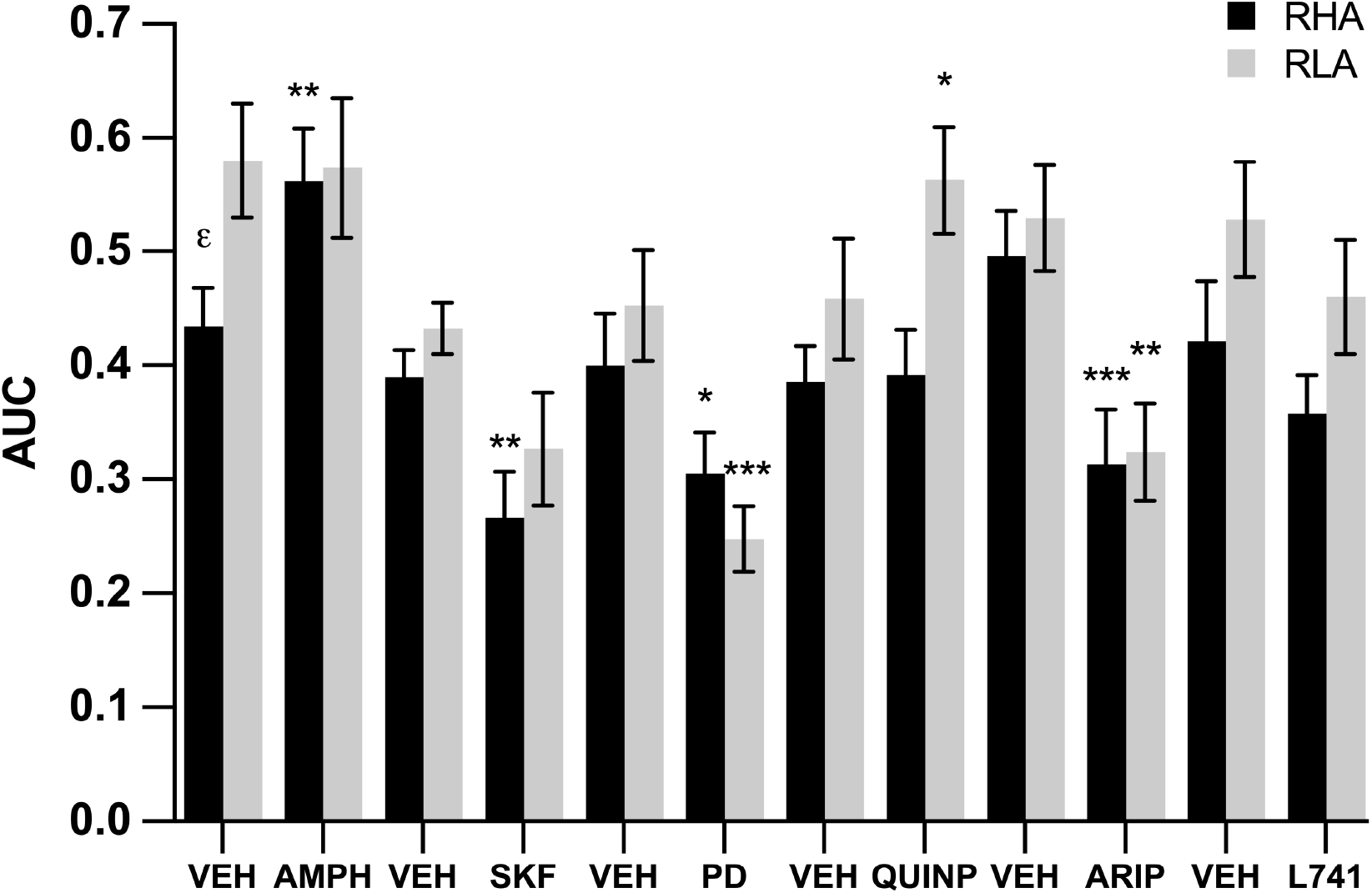
Dopaminergic modulation of AUC in RHA (n= 11-12) and RLA rats. Pharmacological manipulations were performed with i.p. injections of: vehicle (VEH), amphetamine (AMPH; 1.0 mg/kg; i.p.), SKF81297 (SKF; 0.5 mg/kg; i.p.), PD128907 (0.3 mg/kg; i.p.), quinpirole (QUINP; 0.5 mg/kg; i.p.), aripiprazole (ARIP; 1.0 mg/kg; i.p.) and L741,626 (L741; 2.0 mg/kg; i.p.). Drug presentation was counterbalanced across animals. Data are expressed as mean±SEM. *P<0.05, **P<0.01 and ***P<0.001 significantly different from the respective vehicle; εP<0.05 significantly different from RLA vehicle using a two-way mixed factorial ANOVA.

## 4. Discussion

To our knowledge, this is the first study to concurrently measure the relationships between the three facets of impulsivity using a within-subject design in rodents, and to compare the effect of DA drugs across different tests of impulsive behavior considering baseline levels of impulsivity. We found a positive correlation between risk and delay decision-making, indicating that both constructs of impulsive choice are related. Interestingly, we showed that impulsive action positively correlated with risky decision-making but not with delay discounting, suggesting partial overlap between impulsive action and choice. Moreover, the three facets of impulsivity were negatively correlated with speed of decision-making, suggesting that each construct is related to insufficient deliberation before making a decision. Besides, RHA rats showed a more impulsive phenotype in the three facets of impulsivity compared to RLA rats, supporting the view that different impulsive behaviors can be comorbid (Barrus et al., 2015). Our study also indicated that the acute effects of DA drugs differ across dimensions of impulsivity and can produce contrasting outcomes depending on baseline levels of impulsivity. While D_2/3_R agonism decreased impulsive action in high impulsive RHAs, it increased risky decision-making and decreased delay-related impulsive choice in low impulsive RLAs. This indicates that, not only D_2/3_R agonism has divergent effects on impulsive action and impulsive choice, but also that its effects are dependent on baseline levels of impulsivity. Besides, amphetamine effects on impulsive action were independent of baseline levels of impulsivity, while it decreased delay-related impulsive choice in high impulsive RHAs only. Similarly, the effect of D_1_R stimulation depended on baseline levels of impulsivity, increasing delay-related impulsive choice in high impulsive RHAs but not in low impulsive RLAs. These findings indicate only partial overlapping neurochemical substrates of the three constructs of impulsivity and further illustrate that baseline levels of impulsive behavior are key factors to consider when examining drug effects.

Compared to RLAs, RHA rats displayed greater impulsive action, riskier decision-making, and higher delay-related impulsive choice, strengthening the view that the three facets of impulsivity can coexist in the same individual. These results expand previous independent studies showing that RHAs or rats selected for high impulsivity on the 5-CSRTT display higher risky decision-making (Barrus et al., 2015) and greater delay-related impulsive choice (Moreno et al., 2010; Robinson et al., 2009). Thus, our study demonstrates the comorbid nature of impulsivity in a normal population, extending previous human studies in impulsive-related disorders such as pathological gambling (PG) and substance use disorders (SUD), where multiple forms of impulsivity co-occur (Mestre-Bach et al., 2020; Michalczuk et al., 2011; Rash et al., 2016). Interestingly, we found that impulsive action was related to some but not all sub-dimensions of impulsive choice. Indeed, impulsive action positively correlated with risky decision-making, which is consistent with other studies (Barrus et al., 2015; Higgins et al., 2018; Tremblay et al., 2021), but not with delay-related impulsive choice, which is also consistent with previous studies (Broos et al., 2012; Winstanley et al., 2004). Still, and although not correlated, impulsive action and delay discounting were higher in RHAs vs. RLA rats, suggesting that these two dimensions of impulsivity may still have common sources of variability. Motor impulsivity and delay discounting tasks assess “waiting impulsivity” but on different timescales (review in Dalley and Ersche, 2019). While impulsive action depends on the instant self-restraint, delay discounting requires subjective decisions over longer time frames to mainly reflect on consequences before the impulsive act (review in Dalley and Ersche, 2019). Collectively, our findings suggest that impulsive action and delay-related impulsive choice represent different components of impulsivity with overlapping mechanisms, making it possible for the two facets of impulsivity to coexist within the same individual without being directly correlated. Additionally, we found that that the two dimensions of impulsive choice, risky decision-making and delay-related impulsivity, were correlated, which is consistent with some (Kirkpatrick et al., 2014), but not a majority of studies addressing this using a within-subject design in rodents (Freels et al., 2020; Gabriel et al., 2019; Mitchell et al., 2012; Shimp et al., 2015; Simon et al., 2009). Given that choices involving delay and choices involving risk are two aspects of a single cognitive process (Benzion et al., 1989), the lack of association between these two sub-dimensions of impulsive choice in earlier studies is puzzling. One might have expected that the two sub-dimensions of impulsive choice are related as both risky decision-making and delay discounting share common features of decision-making in which “costs” –risk of punishment and delay to reward delivery– are interconnected forms of impulsivity (Talmi and Pine, 2012). Indeed, in humans, there is a few studies showing that the two facets of impulsivity are related (Grecucci et al., 2014; Mestre-Bach et al., 2020; Stea et al., 2011). Therefore, finding the same correlation here in rodents supports the validity of using animal models to improve our understanding of impulsivity. Conflicting results with past evidence may stem from the narrow range or small inter-individual variations in impulsivity found in inbred rat strains, which could have hampered the detection of possible relationships between impulsivity constructs. Consequently, by expanding the range of impulsivity using a cohort of animals with various levels of innate impulsivity may have allowed us to reveal relationships between different measures of impulsivity.

Interestingly, RHA rats were faster to make a choice than RLAs in both the rGT and the DDT, suggesting that the impulsive phenotype in RHAs could partially result from insufficient reflection. Reflection refers to the tendency to evaluate information before making a decision (Kagan, 1966) and the lack of it might represent a potential mechanism by which decision-making could be biased toward suboptimal choices. Furthermore, the association found between the three facets of impulsivity and choice latency suggests that faster speed of responding was predictive of higher impulsive behavior. Similarly, Barrus et al. (2015) found a negative correlation between motor impulsivity and speed of decision-making. In contrast, RHA rats displayed slower latency to collect the reward than RLA rats in both the rGT and the DDT, although this effect did not reach statistical significance in the rGT, suggesting that RHA rats might have less motivation for food. However, decreased motivation for food in RHA rats is unlikely since RHA rats display higher preference and intake of palatable food (Giorgi et al., 1999; Timgren, 1975). Therefore, the slower reward collection latency in RHA rats could indicate that animals were more interested in the nose-poke hole (i.e., conditioned stimulus), rather than in the pellet reward itself. These results reveal a sign-tracking behavior (preference for the stimulus predictive of reward over the reward itself) in RHA rats, which has been suggested to promote suboptimal behavior in decision-making tasks (Chow et al., 2017; Swintosky et al., 2021). Consistent with this, delay discounting was negatively correlated with collect latency, indicating that motivation for food was predictive of low delay-related impulsive choice.

Replicating and extending previous studies performed in different cohorts of animals (Baarendse et al., 2013; Barrus et al., 2015; Swintosky et al., 2021; Zeeb et al., 2016), we found that AMPH produced contrasting effects across different dimensions of impulsivity when using a within-subject design. It increased impulsive action but had no effect on risky decision-making and we further showed that both these effects were irrespective of baseline levels of impulsivity. Although, this is the first demonstration that AMPH effect on risky decision-making is unaffected by baseline levels of impulsivity, previous reports have yielded mixed results on the effects of psychostimulants on impulsive action in rats, with some showing baseline-dependent effects (Caprioli et al., 2015; Eagle et al., 2007; Feola et al., 2000) and others showing similar effects in both behavioral phenotypes (Barlow et al., 2018a; Fernando et al., 2012). On the other hand, AMPH reduced delay discounting but only in animals with high baseline levels of impulsivity, a result consistent with previous studies (Barbelivien et al., 2008; Higgins et al., 2021), and that may explain some of the inconsistent findings reported on this topic (Baarendse and Vanderschuren, 2012; Broos et al., 2012; Isherwood et al., 2017; Koffarnus et al., 2011). The effects of AMPH on impulsivity thus depends on the specific dimension of impulsivity measured and, at least partly, on baseline levels of impulsivity. Besides acting on the DA transporter and increasing dopaminergic tone, AMPH also affects the serotonin and noradrenaline systems (Kuczenski and Segal, 1997), which also have important roles in modulating impulsivity (review in Dalley and Roiser, 2012). However, current evidence indicates that the pro-dopaminergic action of AMPH underly its differential effects on the different sub-components of impulsivity. Indeed, DA receptor blockade, particularly at the D_1_R and D_2/3_R, can prevent or attenuate AMPH-induced increase in inhibitory control (Pattij et al., 2007; van Gaalen et al., 2006a; van Gaalen et al., 2009) as well as AMPH-induced decrease in delay discounting (van Gaalen et al., 2006b). Moreover, the selective DA transporter inhibitor GBR12909 reproduces AMPH effects on impulsive behaviors, increasing impulsive action in the 5-CSRTT (Fernando et al., 2012) and rGT (Baarendse et al., 2013), but decreasing delay discounting in the DDT (van Gaalen et al., 2006b), while having no effect on risky decision-making in the rGT (Baarendse et al., 2013). On the other hand, increasing noradrenergic or serotonergic tone through selective reuptake inhibitors decrease impulsive action in the 5-CSRTT (Higgins et al., 2021; Humpston et al., 2013), with null effects on delay discounting and risky decision-making (Baarendse et al., 2013; Higgins et al., 2021; Paterson et al., 2012). Intriguingly though, and along with previous observations (review in Winstanley, 2011), AMPH effects on impulsivity could not be recapitulated by stimulating DA receptors. Even more intriguingly, D_1_R, D_2/3_R and D_3_R stimulation had opposite effects to those of AMPH on impulsive action as it decreased rather than increasing premature responding. Along the same line, D_1_R and D_2/3_R agonists increased, whilst AMPH had no effect on risky decision-making. Only D_2/3_R stimulation with quinpirole, but not D_3_R stimulation with PD128907, had an effect alike AMPH on delay discounting. As quinpirole has high affinity for both D_2_R and D_3_R (Malmberg and Mohell, 1995) and PD128907 is more selective for D_3_R than D_2_R (Pugsley et al., 1995), the reducing effect of AMPH on delay-related impulsivity may more selectively involve the DA D_2_R subtype. One explanation for the divergent effects of AMPH and individual DA receptor stimulations on impulsivity is that AMPH effects may result from a balanced contribution, and possibly opposing, of D_1_-like and D_2_-like signaling rather than simply activation of one or the other receptor subtypes.

In agreement with previous studies demonstrating that stimulation of D_1_R (Pekcec et al., 2018; Winstanley et al., 2010b) and D_3_R (Barrus and Winstanley, 2016) decreased premature responding, we further showed that both these effect were independent of baseline levels of impulsivity. In contrast, the effect of D_2/3_R stimulation was dependent on baseline levels of impulsive action, with premature responding being decreased in high impulsive RHAs but not in low impulsive RLAs. Consistent with previous studies though (Swintosky et al., 2021; Winstanley et al., 2010b; Zeeb et al., 2009), D_2/3_R stimulation with quinpirole also produced strong other effects, increasing the rate of omissions and the latencies to respond and to collect the reward. It is worth noting that the vast majority of studies to date on impulsivity have used low doses of quinpirole (typically ≤ 0.125 mg/kg). However, at low doses, quinpirole produces a general reduction in motor output (Eilam and Szechtman, 1989; Tournier et al., 2013), an effect that has been attributed to the activation of presynaptic D_2/3_ autoreceptors (White and Wang, 1984) and may explain poor performances in rapid-response impulsivity tasks. In the present study, we used a higher dose of quinpirole (i.e. 0.5 mg/kg), which, in contrast, produces a general activation in motor output (Eilam and Szechtman, 1989; Tournier et al., 2013) likely through stimulation of postsynaptic D_2/3_R in striatum (Kling-Petersen et al., 1995). Surprisingly, even at a dose increasing motor function, quinpirole also decreased performances in the rGT, an effect that may reflect a decreased motivation leading to a disengagement from the task. Consistent with previous studies (Di Ciano et al., 2015; van Gaalen et al., 2009), D_2_R antagonism did not affect premature responding and, this effect was independent of the baseline levels of impulsivity. In contrast, partial D_2/3_R agonism with aripiprazole decreased premature responding irrespective of baseline levels of impulsive action, as previously shown (Besson et al., 2010). However, contrasting with this latter study, aripiprazole did not induce a general motor slowing, in that the percentage of omissions as well as all response latencies were not affected, thus suggesting a specific effect of the drug on premature responding. Aripiprazole is a DA stabilizer, which either stimulate or inhibit DA-related behaviors depending upon the prevailing DAergic tone (DeLeon et al., 2004). Given that RHA and RLA rats show divergent functional properties of their DA neurotransmission system, with RHA rats displaying lower striatal D_2/3_R availabilities and higher AMPH-induced DA release in striatum than RLA rats (Belles et al., 2020; Tournier et al., 2013), we postulated that aripiprazole may have different effects on impulsivity in both lines. At the dose used here (i.e., 1 mg/kg), aripiprazole produces 60% occupancy of the D_2/3_R with weak occupancy of the 5HT_2A_receptor (Natesan et al., 2006). However, aripiprazole similarly decreased impulsive action in RHA and RLA rats, suggesting its effects were not dependent of individual baseline levels of impulsive action but also not dependent on the status of DA functioning. Alternatively, and together with the lack of effect of D_2_R blockade, these results suggest that the reducing effects of aripiprazole on impulsive action may not be mediated by its D_2/3_R partial agonistic properties. On the other hand, aripiprazole also has 5-HT_1A_antagonistic properties (review in Casey and Canal, 2017; Tuplin and Holahan, 2017), that may also account for the decreased impulsive action as 5-HT_1A_antagonism has been shown to reduce premature responding (Ohmura et al., 2013; Quarta et al., 2007).

Contrasting with its enhancing effects on premature responding, amphetamine did not affect risky decision-making in either RHA or RLA rats. Besides, risky decision-making seemed to be affected by stimulation of D_1_R and D_2/3_R but not D_3_R. While in previous studies D_1_R agonism increased risky decision-making (St Onge and Floresco, 2009; Zeeb et al., 2009), we further observed that this effect was regardless of the baseline levels of impulsivity. In contrast, D_2/3_R stimulation produced baseline-dependent effects, as it increased risky decision-making in low impulsive RLAs only. Such a baseline-dependent effect might explain some of the inconsistencies found in the literature regarding the effect of D_2/3_R agonist on risky decision-making (Georgiou et al., 2018; Simon et al., 2011; Zeeb et al., 2009). In contrast, D_2_R blockade and aripiprazole had no effect on risky decision-making, suggesting that D_2_R and 5-HT_1A_R antagonisms are likely minimally involved in risky decision-making, which is supported by previous data (Blaes et al., 2018; Di Ciano et al., 2015).

In the DDT, the effect of AMPH was dependent on baseline levels of impulsivity, decreasing delay-related impulsive choice in high impulsive RHA rats only. Other studies have found that psychostimulants’ effects on the DDT are dependent upon baseline levels of impulsive choice (Bickel et al., 2016; Higgins et al., 2021; Winstanley et al., 2003). Our results further indicates that the effects of AMPH on delay discounting are also dependent on the baseline levels of impulsive action. Besides, whereas D_1_R and D_3_R stimulation increased delay discounting irrespective of baseline levels of impulsivity, D_2/3_R agonism decreased it in low impulsive RLAs only. Thus, stimulation of D_1_R and D_3_R on one hand and stimulation of D_2/3_R on the other hand had dissociable and opposite effects that depend in part on baseline levels of impulsivity. Moreover, while D_2_R blockade had no effect in RHA and RLA rats, partial D_2/3_R agonism with aripiprazole increased delay-related impulsive choice regardless of baseline levels of impulsivity.

The dissociable effects of systemic administration of DA agents across dimensions of impulsivity have also been observed in studies using tools with greater specificity and precision of the neural circuits. The nucleus accumbens (NAc) controls different facets of impulsivity (Basar et al., 2010) but its core and shell sub-regions have been proposed to have distinct roles in modulating impulsive behaviors. Thus, optogenetic stimulation of the ventral striatum (VTA)-NAc shell pathway increased impulsive action (Flores-Dourojeanni et al., 2021), while stimulation of VTA-NAc core pathway decreased delay-related impulsive choice (Saddoris et al., 2015). An involvement of the NAc in risky decision-making is also supported by ex- and in-vivo DA receptor expression in NAc shell but not core (Freels et al., 2020; Simon et al., 2011). In contrast, reversible inactivation of infralimbic cortex by the GABA_A_ receptor agonist muscimol did not affect delay discounting (Feja and Koch, 2014) which seems to recruit more the dorsal area of the prefrontal cortex (PFC), namely prelimbic cortex (Sackett et al., 2019). However, optogenetic studies have shown that the infralimbic plays a more prominent role in modulating impulsive action (Feja and Koch, 2014; Hardung et al., 2017) and risky decision-making (Zeeb et al., 2015). Thus, anatomical selectivity of subregions in the NAc or PFC may be one possible explanation for the dissociable behavioral effects among the three dimensions of impulsivity observed in the present study.

## Supporting information

SupInfo

## Declaration of interest

The authors declare that there is not personal financial conflict of interest.

## Role of the funding source

This study was supported by the Swiss National Science Foundation (SNF; grant number: 31003A_179373). SNF had no further role in study design; collection, analysis, or interpretation of data; writing of the report; or the decision to submit the paper for publication.

## Acknowledgments

This study was supported by the Swiss National Science Foundation (SNF; grant number: 31003A_179373).

## Notes

### Competing Interest Statement

The authors have declared no competing interest.

