## Supplementary material for "Concurrent measures of impulsive action and choice are partially related and differentially modulated by dopamine D_1_- and D_2_-like receptors in a rat model of impulsivity": SupInfo

**Pharmacology Biochemistry and Behavior**

**This file includes:**

**Supplementary methods**
**Table S1 to S4**
**Supplementary references**

**Supplementary methods**

### Rat Gambling Task (rGT)

Rats were trained on the rGT as previously described (Zeeb et al., 2009).

*Training.* Animals were habituated to the food pellet rewards and the operant chamber for two 30-min sessions, in which all lights of the chambers were turned on and pellets were placed in the response holes and food magazine. Animals were then trained to make a nose poke response into an illuminated hole first within 30 sec (training phase 1), then 20 sec (training Phase 2) and 10 sec (training Phase 3) for a single reward. Once rats completed these sessions with at least ≥50 correct trials, ≥80% accuracy and <30% omissions, animals received seven sessions of a force-choice version of the rGT, in which only one option was presented per trial, to ensure equal experience with all the four reinforcement contingencies and avoid preferences toward a particular hole.

*rGT.* Training was conducted for about 25 daily sessions (5 days per week). Each session had a duration of 30 min or 100 trials. At the beginning of each session, the house light was illuminated, and the trial was initiated when the animal made a nose-poke response into the illuminated food tray. After an inter-trial interval (ITI) of 5 sec, the four stimulus lights in holes 1, 2, 4 and 5 (termed P1, P2, P3 and P4 respectively) were illuminated for a period of 10 sec and the animal had to choose any one of these options by making a nose-poke response into the corresponding hole. A nose-poke into any of these holes turned on the food tray light and the corresponding number of pellets were delivered in the receptacle (1, 2, 3 or 4, respectively) each with a defined probability (90%, 80%, 50% and 40% respectively). Collection of this reward initiated the next trial. However, if the trial was not rewarded, the stimulus light within the hole that the animal chose flashed at 0.5 Hz for the duration of the corresponding punishing timeout (TO) period (5, 10, 30 or 40 sec, respectively). Missed trials (i.e., omissions) were also recorded and all stimulus lights were extinguished, and the food tray light turned on, allowing the animal to start another trial. Responses made before the onset of the visual stimulus (i.e., during the ITI) were considered as premature or impulsive and resulted in a 5-sec TO period. Following this TO, the food tray was illuminated, and the animal could initiate the next trial. Two versions of the rGT were used which differed only in the spatial location of the options and were counterbalanced across all animals. In a version A, the order of the options presented from left to right was P1 (one-pellet option), P4 (four-pellet option), P2 (two-pellet option) and P3 (three-pellet option). In version B, the order of the option from left to right was P4, P1, P3 and P2. The optimal choice in rGT was P2, as this option is the most rewarded per unit time. The next best option was P1, and the two disadvantageous options were P3 and P4, due to the lower probability of receiving reward and the longer punishing TO periods incurred. Summary of the reward and punishment options in the different versions of the rGT is shown in Table S2. The following rGT variables were analyzed: choice score [(P1+P2)–(P3+P4)*100], a measure of risky decision making, and percentage of premature responses [(#premature responses/#total number of trials initiated)*100] as a measure of motor impulsivity.

### Discounting Delay Task (DDT)

*Training.* Animals were initially exposed to a modified version of the amount-discrimination training (Renee Renda et al., 2018) for magnitude discrimination. Each session had a duration of 72 min and was divided into 6 blocks of 12 trials, constituted of 4 forced-choice trials followed by 6 free-choice trials. Rats were trained to nose-poke in the food tray to trigger the illumination of one (forced-choice trials) or the two (free-choice trials) holes. A nose-poke into the small reinforcer hole turned on the food tray light and one pellet was delivered in the receptacle with no time delay (i.e., 0s). If the choice was the large reinforcer hole the stimulus light switched off, the food tray was illuminated, and 3 pellets were delivered also with no time delay. An adjusting intertrial interval ensured that a new trial started every 72 sec. Sessions ended when all 60 trials were completed or if 72 min elapsed. This training was conducted to ensure that rats were not avoiding responding the three-pellet option during the trial blocks with delay. Rats were required to reach a criterion of 60 completed trials with ≥80% preference of choice of the three-pellet alternative across three consecutive sessions (approx. 10 to 15 sessions).

*DDT.* DDT was conducted for about 35 daily sessions (5 days per week) as previously described (Barlow et al., 2018; Isherwood et al., 2017). Sessions were structured identically to the training sessions, with the exception that the delay to the three-pellet option increased systematically across all blocks from 0 to 2, 4, 8, 16 and 32 sec. Missed trials (i.e., omissions) were also recorded and the trial was directly entering into an inter-trial interval initiated before the next trial. Delay discounting was represented by the Area Under the Curve (AUC), defined as the area under the discounting curve divided by the total area of the discounting graph. AUC was calculated based on Magnard et al. (2018); Myerson et al. (2001). Briefly, delays, and large reinforcer preference were first expressed as the proportion of their maximum value to be comprised between 0 to 1. Then, the resulting discounting curve was subdivided discounting graph into a series of half trapezoids, from 0 to 2, 2 to 4, 4 to 8, 8 to 16 and 16 to 32 sec. The area of each trapezoid is thus equal to the following equation $x2 - x1(y1 + y2)/2$, where $x1$ and $x2$ are successive delays associated with the larger reinforcer preference $y1$ and $y2$, respectively. For delay discounting, smaller AUC values means a steep discounting (high impulsivity), whereas larger AUC values means little to no discounting (low impulsivity).

**Supplementary Tables**

**Table S1.** Details of drugs doses used

| Drug | Drug type | Dose | Pretreatment time |
|---|---|---|---|
| **Amphetamine** | Nonselective dopamine agonist | Vehicle, 1.0 mg/kg | 90 minutes |
| **SKF81297** | Dopamine D1 agonist | Vehicle, 0.5 mg/kg | 30 minutes |
| **Quinpirole** | Dopamine D2/3 agonist | Vehicle, 0.5 mg/kg | 90 minutes |
| **Aripiprazole** | Dopamine partial D2/3 agonist | Vehicle, 1.0 mg/kg | 20 minutes |
| **L-741,626** | Dopamine D2 antagonist | Vehicle, 2.0 mg/kg | 60 minutes |
| **PD128907** | Dopamine D3 agonist | Vehicle, 0.3 mg/kg | 20 minutes |

**Table S2.** Reward and punishment received for the various response options in the rGT (Zeeb et al., 2009)

| **Version A** | Hole 1 | Hole 4 | Hole 5 | Hole 2 |
|---|---|---|---|---|
| **Version B** | Hole 2 | Hole 5 | Hole 4 | Hole 1 |
| **Choice** | P1 | P2 | P3 | P4 |
| **Number of pellets rewarded** | 1 | 2 | 3 | 4 |
| **Change of a win trial** | 90% | 80% | 50% | 40% |
| **Punishment duration (s)** | 5 | 10 | 30 | 40 |
| **Rewards possible** | 295 | 411 | 135 | 99 |

**Table S3.** Total number of trials, percentage of omissions, choice and collect latencies of the different pharmacological manipulations in RHA and RLA rats on the rGT.

| | Line | Veh | AMPH | Veh | SKF | Veh | PD | Veh | QUINP | Veh | ARIP | Veh | L741 |
|---|---|---|---|---|---|---|---|---|---|---|---|---|---|
| **Omission (%)** | **RHA** | 5 ± 0.9 | 2 ± 1.1 | 7 ± 1.3 | 9 ± 1.4 | 8 ± 1.2 | 21 ± 3.7** | 6 ± 1.1 | 31 ± 3.0*** | 5 ± 1.3 | 7 ± 1.2 | 4 ± 1.2 | 7 ± 1.7 |
| | **RLA** | 22 ± 2.3 | 15 ± 7.2 | 26 ± 2.3 | 36 ± 5.9 | 25 ± 2.2 | 32 ± 3.1 | 25 ± 2.2 | 55 ± 10.0* | 25 ± 2.6 | 9 ± 3.4 | 20 ± 2.2 | 25 ± 2.6 |
| **Choice latency (s)** | **RHA** | 2 ± 0.3 | 1.8 ±0.6 | 2.1 ± 0.2 | 2.3 ± 0.2 | 2.2 ± 0.3 | 2.7 ± 0.3 | 1.7 ± 0.2 | 2.8 ± 0.2** | 1.6 ± 0.2 | 2 ± 0.2 | 1.6 ± 0.2 | 1.8 ± 0.2 |
| | **RLA** | 3.9 ± 1.1 | 1.6 ± 0.3 | 2.6 ± 0.1 | 2.5 ± 0.3 | 2.5 ± 0.2 | 2.6 ± 0.2 | 2.5 ± 0.2 | 2.5 ± 0.3 | 2.4 ± 0.1 | 2.7 ± 0.2 | 2.6 ± 0.1 | 2.5 ± 0.3 |
| **Collect latency (s)** | **RHA** | 1 ± 0.1 | 1.6 ± 0.4 | 1.1 ± 0.0 | 1.2 ± 0.0 | 1 ± 0.1 | 1.7 ± 0.3* | 1.1 ± 0.1 | 1.7 ± 0.2* | 1.1 ± 0.1 | 1.1. ± 0.1 | 1.2 ± 0.1 | 1 ± 0.1 |
| | **RLA** | 1 ± 0.1 | 0.8 ± 0.1 | 1 ± 0.0 | 1.1 ± 0.2 | 1.2 ± 0.2 | 1.3 ± 0.2 | 1.3 ± 0.4 | 2.2 ± 0.3** | 1 ± 0.0 | 1 ± 0.1 | 1 ± 0.1 | 1 ± 0.1 |

Data are mean ± SEM. *$P<0.05$, **$P<0.01$ and ***$P<0.001$ using paired Student's t test.

**Table S4.** Choice and collect latencies of the different pharmacological manipulations in RHA and RLA rats on the DDT.

| | Line | Veh | AMPH | Veh | SKF | Veh | PD | Veh | QUINP | Veh | ARIP | Veh | L741 |
|---|---|---|---|---|---|---|---|---|---|---|---|---|---|
| **Choice latency (s)** | **RHA** | 1.3 ± 0.1 | 1.4 ± 0.2 | 2 ± 0.3 | 2.6 ± 0.4 | 1.7 ± 0.2 | 2.3 ± 0.2 | 1.3 ± 0.1 | 3.6 ± 0.8** | 1.6 ± 0.2 | 1.8 ± 0.2 | 1.6 ± 0.2 | 1.8 ± 0.3 |
| | **RLA** | 2.5 ± 0.3 | 1.7 ± 0.1* | 2.5 ± 0.3 | 2.6 ± 0.3 | 2.5 ± 0.3 | 2.7 ± 0.2 | 2.9 ± 0.3 | 4.7 ± 0.6** | 3.4 ± 0.7 | 3.8 ± 0.6 | 2.6 ± 0.4 | 2.7 ± 0.4 |
| **Collect latency (s)** | **RHA** | 0.5 ± 0.0 | 0.5 ± 0.1 | 0.4 ± 0.0 | 0.6 ± 0.0* | 0.5 ± 0.0 | 0.7 ± 0.1** | 0.5 ± 0.0 | 0.7 ± 0.1** | 0.4 ± 0.0 | 0.6 ± 0.0* | 0.5 ± 0.0 | 0.5 ± 0.0 |
| | **RLA** | 0.3 ± 0.0 | 0.5 ± 0.1* | 0.5 ± 0.0 | 0.5 ± 0.0 | 0.4 ± 0.0 | 0.7 ± 0.1*** | 0.4 ± 0.0 | 0.9 ± 0.1** | 0.5 ± 0.1 | 0.5 ± 0.1 | 0.5 ± 0.1 | 0.4 ± 0.1 |

Data are mean ± SEM. *$P$<0.05, **$P$<0.01 and ***$P$<0.001 using paired Student's t test.
